# Machine learning for optimal growth temperature prediction of prokaryotes using amino acid descriptors

**DOI:** 10.1101/2025.03.03.640802

**Authors:** Sophie Colette, Jaldert François, Bart De Moor, Vera van Noort

**Affiliations:** KU Leuven, Centre of Microbial and Plant Genetics, Faculty of Bioscience Engineering, Leuven, Belgium; KU Leuven, Department of Electrical Engineering (ESAT), Leuven 3001, Belgium; Leiden University, Institute of Biology Leiden (IBL), Leiden 2333 BE, The Netherlands

## Abstract

**Motivation:** The optimal growth temperature (OGT) of organisms is valuable in bioprospecting enzymes that work under extreme conditions. Existing OGT prediction models achieve high accuracy, but mainly capture trends of overrepresented groups in the training set including organisms that thrive at moderate temperatures and those from well-described taxa.

**Results:** In this study, we incorporated weighted scoring and phylogenetic splits to improve the generalizability of the prediction models. We first built a new growth temperature dataset comprising more than 21,000 species distributed over all three domains of life, with special attention to include OGT and extreme temperature data. We then trained machine learning models on the OGT data of 6,401 prokaryotes using proteome-averaged amino acid descriptors. The best-performing model was the multilayer perceptron with a cross-validated RMSE of 5.07°C (*±* 0.24) and an R^2^ of 0.89 (*±* 0.04). The most important proteome features were related to backbone flexibility, charged residues, as well as surface accessibility.

**Availability and Implementation:** The MLP model is integrated in the command line tool OGTFinder and available under MIT license at: https://github.com/SC-Git1/OGTFinder.

## 1 Introduction

Bioprospecting identifies novel molecules, enzymes, and microbes relevant to biological and biochemical research and industrial applications. Of particular interest for the bioprospecting of microbes and enzymes are process parameters such as temperature. Temperature affects the internal energy of the cell, and temperature directly impacts every cellular component (Ramón et al., 2023; Wang et al., 2015). For example, temperature has a large influence on protein stability, with higher temperatures leading to denaturation, protein aggregation, and eventual loss of function, while low temperatures lead to reduced enzyme kinetics and eventual inactivation (Plaza Del Pino et al., 2000; D’Amico et al., 2002). Each organism is adapted to a specific temperature range (Corkrey et al., 2016). At the optimal growth temperature (OGT), an organism achieves its maximum growth rate. According to their OGT, organisms are broadly classified into psychrophiles (OGT *<* 15°C), mesophiles (15°C *<* OGT *<* 45°C), thermophiles (45°C *<* OGT *<* 80°C) and hyperthermophiles (OGT *>* 80°C), although these temperature boundaries are not fixed (Jensen et al., 2012).

OGT annotation is valuable for investigating general trends in thermal adaptation or specific trends through comparative analysis. OGT has been found to correlate with properties of the genome (Kawashima et al., 2000; Hu et al., 2022; Zeldovich et al., 2007; Sabath et al., 2013), rRNA and tRNA (Galtier and Lobry, 1997; Hurst and Merchant, 2001; Khachane et al., 2005), codon usage (Lobry and Chessel, 2003; Sauer and Wang, 2019), proteome (Zeldovich et al., 2007; Kurokawa et al., 2023; Sauer and Wang, 2019; Tekaia and Yeramian, 2006; Villain et al., 2022), and metabolic network (Weber Zendrera et al., 2019; Farrell et al., 2025). Furthermore, OGT correlates with enzyme optimal catalytic temperatures (r = 0.76; Wang et al. 2024), and current prediction models of these temperatures rely heavily on OGT (Li et al., 2019; Gado et al., 2020; Li et al., 2022). Thus, by estimating optimal enzyme catalytic temperatures, OGT indirectly informs species selection for bioprospecting enzymes.

Experimentally determining OGT is a laborious and time-consuming process, as it requires incubation and growth analysis at various temperatures (Sauer and Wang, 2019; Yang et al., 2024). This method is limited to culturable organisms, which are biased towards certain phyla and only account for a small fraction (roughly 3%) of all prokaryotes (Lewis et al., 2021). Initial approximation of growth temperature from environmental data of metagenomics can be inaccurate due to local temperature gradients (Khachane et al., 2005), or influx of micro-organisms (Kurokawa et al., 2023). Furthermore, organisms can tolerate temperatures far from their OGT, such as facultative thermophiles whose OGT is larger than 45°C, but can also grow at moderate temperatures (Bergey, 1919). Similarly, psychrotrophs are cold-tolerant but have an OGT above 15°C (Morita, 1975). OGT predictions can thus (1) enhance the culturability of organisms and (2) avoid the high cost and other challenges associated with experimental determination of optimal growth conditions.

Due to its importance, predicting OGT has become a popular research topic with a continuous release of dedicated databases, such as TEMPURA (Sato et al., 2020) and ThermoBase (DiGiacomo et al., 2022), and the development of several OGT prediction models based on features of the genome (Wang et al., 2022), tRNA (Cimen et al., 2020), proteome, (Li et al., 2019; Kurokawa et al., 2023; Barnum et al., 2024), or a combination thereof (Jensen et al., 2012; Sauer and Wang, 2019). Despite the progress, current OGT prediction models face several limitations. One key issue involves the phylogenetic signal in OGT data (Hu et al., 2022). Random train-test splits may cause models to overfit on taxonomic patterns, resulting in overly optimistic test performances. This limits their ability to generalize to distantly related organisms and gain biological insights (Cimen et al., 2020). Moreover, mesophiles are often overrepresented in training data, reducing model performance at the extremes, especially at low temperatures (Sauer and Wang, 2019; Li et al., 2019). While some studies have attempted to address this bias through undersampling of mesophiles (Wang et al., 2022; Cimen et al., 2020), others did not (Li et al., 2019; Sauer and Wang, 2019). Finally, the lack of extensive model explanation for most OGT prediction models has limited the identification of the most predictive features and their coupling back to underlying biological mechanisms.

Here, we developed an OGT prediction model based on physicochemical amino acid properties (Osorio et al., 2015) from predicted proteomes. Our approach includes phylogenetically informed data splits, weighted RMSE for better predictions in extremophiles, and the SHAP framework for model interpretation (Lundberg and Lee, 2017). The model is integrated in a command line tool OGTFinder available at https://github.com/SC-Git1/OGTFinder.

## 2 Materials and Methods

### 2.1 Optimal growth temperature database construction

#### 2.1.1 Data retrieval

Optimal growth temperature (OGT) data was downloaded from four databases (ThermoBase (DiGiacomo et al., 2022), TEMPURA (Sato et al., 2020), aciDB (Neira et al., 2020), MediaDB (Richards et al., 2014)) and the supplementary material of Lyubetsky et al. (2020). In addition, cultivation temperature data was obtained from three culture collection centers (CCUG ^1^, CECT ^2^, NIES ^3^) through Python web scraping. Finally, temperature records were extracted from the bacterial metadatabase BacDive using the API (Reimer et al., 2022). The data extraction was performed from February to April 2023.

#### 2.1.2 Taxonomy mapping

Extracted records were mapped to a unique taxonomic identifier, the NCBI TaxId, based on organism name(s) and with the help of the NCBI datasets Python API and the Taxonomy browser. If no matching TaxId could be found, manual annotation was attempted by looking up the cross-referencing culture collection numbers, or through other metadata. The preferred TaxId rank was “strain”. Records that could only be identified down to the genus level were removed. Any out-of-use TaxIds were replaced.

#### 2.1.3 Temperature data cleaning

In the case of BacDive, records were of type minimal, maximal, optimum or growth. Minimal and maximal temperatures were removed as these correspond to conditions of no growth. For the remaining databases, all records were uniformly annotated as “optimum” or, if no indication of optimality was present, “growth” for cultivation temperature. Temperature ranges were averaged. Two records of *Levilactobacillus acidifarinae* and *Levilactobacillus zymae*, both with an OGT of 7,5°C in BacDive, were excluded because the annotations could not be verified against the original paper.

#### 2.1.4 Genome extraction and annotation

All prokaryotic TaxIds were mapped to their reference genome or, if unavailable, all their genomes in NCBI using the NCBI datasets command line tool. The species level was used if no genomes were available at the lowest taxonomic level. High-quality genomes were selected based on 1) the average nucleotide identity (ANI) report, and 2) the assembly level. The ANI report contains the result of an ANI-based analysis that evaluates whether the assigned taxonomy of an assembly is correct. All genome assemblies with a ‘taxonomy-check-status’ of “inconclusive” or “failed” were removed, leaving only unassessed genomes and genomes with a correct taxonomic identity. Next, for each TaxId, only assemblies with the highest assembly level were kept following the order ‘Complete Genome *>* Chromosome *>* Scaffold *>* Contig’. The 10,652 assemblies left after filtering were downloaded using the NCBI Datasets command line tool and annotated with Prokka (version 1.14.6, Seemann 2014) using the default installation. The taxonomic domain and genus names were provided.

### 2.2 Machine learning

#### 2.2.1 Predictor

While in the past, OGT prediction models were trained on cultivation temperatures (Li et al., 2019; Sauer and Wang, 2019), most recent models have been trained on OGT (Wang et al., 2022; Barnum et al., 2024). In this work, we only used the OGT data for modelling. Still, the cultivation temperatures serve two purposes: (1) to assess the difference in data quality between cultivation temperature and OGT, and (2) to provide growth conditions for more microorganisms, including extremophiles, for further studies.

#### 2.2.2 Features

Existing OGT prediction models based on proteome information were trained on amino acid or dipeptide composition (Sauer and Wang 2019; Li et al. 2019; Barnum et al. 2024). In this work, we used 58 descriptors that capture key amino acid characteristics derived from dimensionality reduction on physico-chemical and/or structural properties. The hypothesis is that the additional information in the descriptors, compared to merely composition, can further improve the model and facilitate the biological interpretation. The selected amino acid descriptors belong to the descriptor sets z-Scales (Sandberg et al., 1998), T-scale (Tian et al., 2007), ST-scale (Yang et al., 2010), MS-WHIM (Zaliani and Gancia, 1999), VHSE (Mei et al., 2005) Kidera (Kidera et al., 1985), Cruciani (Cruciani et al., 2004), Blosum (Georgiev, 2009) and Fasgai (Liang and Li, 2007). The descriptors were extracted with the R package “peptides” (version 2.4.4, Osorio et al. 2015) and averaged over the entire proteome. In addition, taxonomic domain information was coded into a dummy variable.

#### 2.2.3 Dataset composition

To remove data imbalance originating from the variable number of records per Taxid, we reduced the machine learning dataset to one observation per TaxId, represented by the mean proteome descriptors and the median OGT. We then performed a phylogenetically informed train-test split (90.0%-10.0%), keeping observations of the same genus together (Barnum et al., 2024). Hyperparameter tuning and performance evaluation were carried out using 5-fold cross-validation on the training set, while the test set was reserved for final model evaluation. Feature normalization was performed within each fold with the function *StandardScaler* from scikit-learn (version 1.5.1, Pedregosa et al. 2011). The binned cross-validation performances were calculated using the combined list of cross-validation predictions over the 5 folds.

### 2.3 Machine learning algorithms

We trained 9 linear models (lasso, ridge, Bayesian ridge, elastic net, LARS, LARS-lasso, Huber, PLS, and stochastic gradient descent) and 5 non-linear models (kernel ridge, Gaussian process, SVR, multilayer perceptron and k-nearest neighbor) using the packages scikit-learn. Hyperparameter tuning was performed in Optuna (version 3.6.1, Akiba et al. 2019) using 5-fold cross-validation over 100 trials with a weighted root-mean-square (RMSE) loss function to address the temperature imbalance in the dataset. The weights were calculated on the entire training set by the following two steps:

For observation *i* with OGT value *x*:

1. Calculate a weight proportional to the inverse of the probability density function:

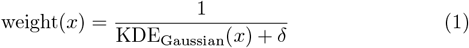

with *δ* = 0.01, and KDE_Gaussian_(*x*) estimated on the training data with the function *scipy*.*stats*.*gaussian kde*
2. Min-max normalize.

### 2.4 Model interpretation

The descriptors most important to the model were identified using the SHAP (SHapley Additive exPlanations) framework. SHAP values, expressed in target units (°C), are estimates of a feature’s marginal contribution to deviations of the predictions from the expected value. SHAP values were calculated with the Python package shap (version 0.46.0, Lundberg and Lee 2017) using the default KernelExplainer and the background set to 10 K-means samples.

#### 2.4.1 Amino acid descriptor interpretation

SHAP values identify influential features, but they rely on the interpretability of those features to provide meaningful biological insights. Some amino acid descriptors applied in this work lack a biological explanation. Therefore, we investigated the correlations of the descriptors with 586 physico- and biochemical amino acid properties from the AAindex (Kawashima et al., 2007), and two additional sources (Koehler et al., 2009; Lins et al., 2003). The indices and their hierarchical categorization were retrieved from the AAanalysis package (Breimann et al., 2024).

### 2.5 Correlation analysis

The Pearson correlation coefficient was used unless stated otherwise. Correlations were classified into strong (|*r*| *≤* 0.7), moderate (0.7 *<* |*r*| *≤* 0.5), and weak (0.5 *<* |*r*| *≤* 0.3). To address the temperature imbalance, the dataset was sorted by OGT and divided into 5 equal-sized bins (n = 1,280), with correlations calculated within each temperature bin. The fifth bin, comprising the top 20% of OGT values, contains temperatures of 37°C or higher.

### 2.6 Phylogenetic signal

The phylogenetic signal in the amino acid descriptors and OGT per taxonomic domain was measured with Pagel’s lambda (Pagel, 1999) on a time-scaled phylogenetic species tree retrieved from TimeTree (Kumar et al., 2022). The tree contained 298 of the 399 archaea and 3,830 of the 5,889 bacteria in our training set. Species with only strain-level information were represented by averaged values. Any zero-length branch lengths were trimmed. The final newick tree was imported with the treeio package (Wang et al., 2020), and Pagel’s lambda was calculated with the function phylosig of the R package phytools (version 2.4-4, Revell 2024).

## 3 Results and Discussion

### 3.1 Dataset

Over 34,000 growth temperature records were collected and assigned to a TaxId, of which 21,881 were cultivation temperatures and 12,864 optimal growth temperatures. The dataset contains mainly bacteria, and almost no eukaryotic data (Table 1). Some data redundancy is expected because of, for example, the requirement for novel species to be deposited in multiple culture collection centers (Rule 30 (3b), Oren et al. 2023) and shared references among OGT databases. The OGT ranges from 2°C to 114°C. The GC content is relatively balanced, with 57% of entries having a GC content of 50% or higher.

**Table 1:** Distribution of TaxIds and species with growth temperature annotation by type and taxonomic domain.

| # | Cultivation |  | Optimal growth |  |
| --- | --- | --- | --- | --- |
|  | TaxIds | Species | TaxIds | Species |
| Archaea | 290 | 269 | 756 | 624 |
| Bacteria | 7,026 | 6,287 | 9,799 | 9,403 |
| Eukaryota | 1,758 | 1,734 | 27 | 27 |

### 3.2 Cultivation versus optimal growth temperature

#### 3.2.1 OGT data is biased towards common culturing temperatures

Earlier OGT prediction models were initially trained on cultivation temperature data (Li et al., 2019; Sauer and Wang, 2019) before switching to OGT (Wang et al., 2022; Barnum et al., 2024). While cultivation temperature refers to any temperature suitable for growth, OGT is the condition that leads to the maximal growth rate. In practice, the OGT is derived from a subset of tested conditions with varying granularity, leading to differences in OGT data quality (Barnum et al., 2024). Kernel density estimations of the median cultivation temperature per species show distinct peaks at common culturing temperatures (CCTs) of Eukaryota and Bacteria (Supplementary Figure S1). In contrast, the kernel density estimation of bacterial OGT shows a reduced bias towards CCTs. The remaining bias is likely due to the choice of evaluation temperatures during the experimental setup. To assess the extent to which cultivation temperatures deviate from optimal growth temperatures, we performed an analysis similar to Wang et al. (2024). Comparison between the median cultivation temperature and the median OGT for the 2,598 species with available data revealed an overall Pearson correlation of 0.90 (Figure 1). 14.2% of the species had a deviation bigger than 5°C, which is about the lower bound of RMSE for current OGT prediction models (Supplementary Figure S2). In 2.6% of the cases (n = 67), the deviation was bigger than 10°C.

**Figure 1:**
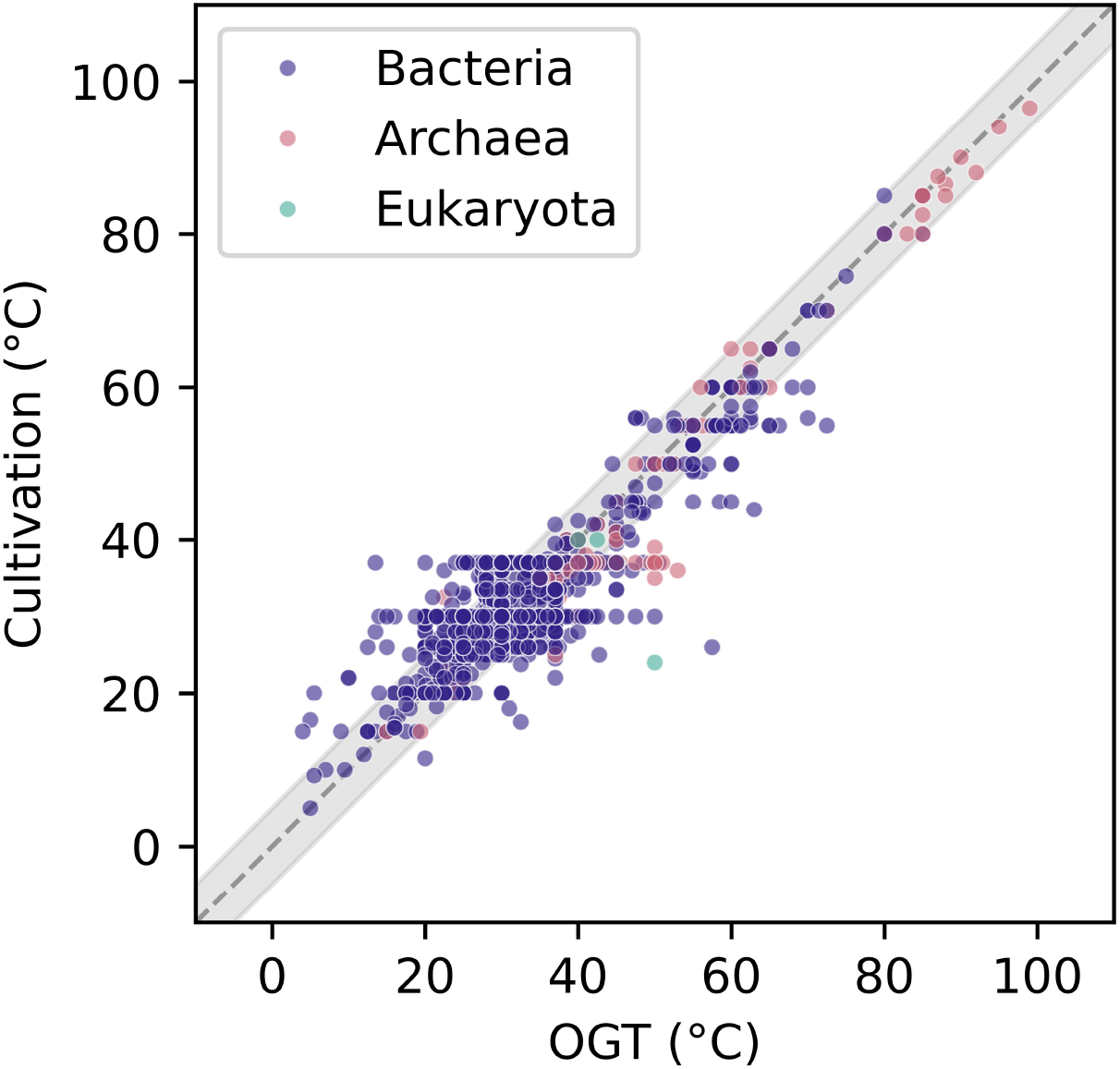
Absolute differences between optimal growth and cultivation temperatures for 2,598 species. Notably, at common culturing temperatures 30°C and 37°C, the range of deviations for cultivation temperatures versus OGT exceed 5°C (highlighted in grey), indicating significantly higher data quality for OGT measurements.

Because of the sometimes large deviations of cultivation temperature from OGT and the negligible amount of eukaryotic OGT data, only prokaryotic optimal growth temperatures were considered for further analysis and model training.

### 3.3 PCA on prokaryotic OGT

#### 3.3.1 Nucleotide bias follows the direction of maximum variance

To identify the directions of highest variance and investigate the clustering patterns, we performed principal component analysis (PCA). The first principal component (PC1) explains 65.68% of the total variance in the features and shows a near-perfect negative correlation (r = -0.97) with GC content (Figure 2 A). Although no genome features were included, the GC content is strongly reflected in the frequencies of amino acids with GC-rich codons: proline (r= 0.95), glycine (r = 0.95), alanine (r = 0.96), and arginine (r = 0.96), as well as AT-rich amino acids lysine (r = -0.96), isoleucine (r = -0.95), glutamine (r = -0.95), tyrosine (−0.90) and phenylalanine (r = -0.88). Methionine (r = -0.47) is the exception among AT-rich amino acids.

**Figure 2:**
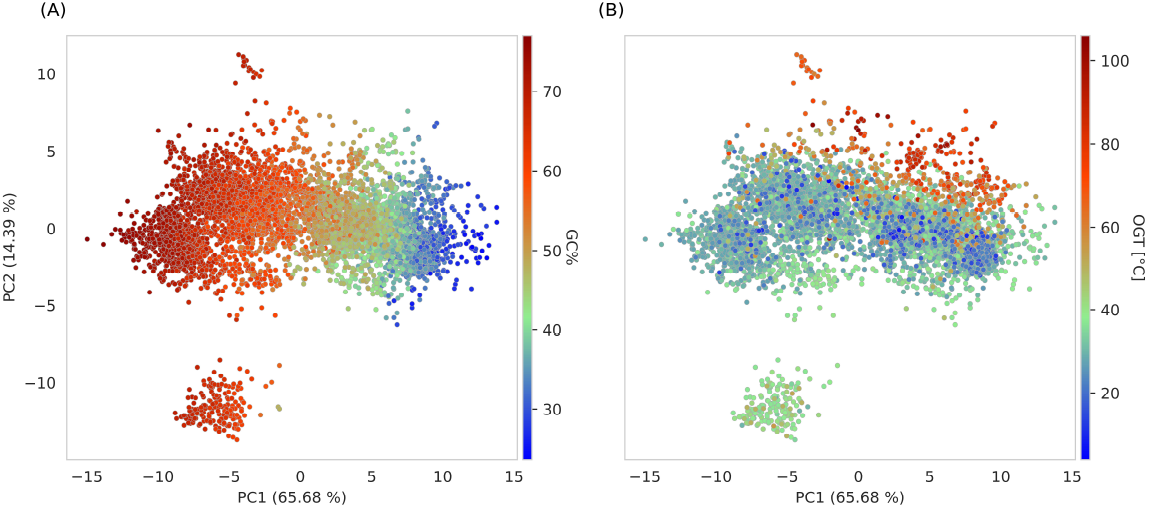
First two principal components (PCs) for the 6,401 TaxIds in the OGT dataset, colored by (A) the average GC percentage, and (B) the median OGT. PC1 follows the direction of decreasing GC percentage. High PC2 values indicate thermophilicity. The outlying observations with low PC2 values belong to the class Halobacteria.

#### 3.3.2 Differences in environmental conditions define PC2

The second principal component, explaining 14.39% of the variance, separates organisms living in extreme conditions, namely, the archaeal class Halobacteria at low values and to a lesser extent, the thermophiles at high values (Figure 2 B). The distinct grouping of Halobacteria is not due to a taxonomic domain split, as other archaea cluster together with bacteria (Supplementary Figure S3, S4).

PC2 correlates strongly negative with aspartate (r = -0.80), moderately negative with threonine (r = -0.59) and moderately positive with the fraction of leucine (r = 0.66). The aspartate fraction combines two trends in extremophilic adaptation. Thermophiles avoid aspartate due to its chemical instability at high temperatures (Villain et al., 2022). In contrast, halophiles prefer negatively charged amino acids as a negative design mechanism promoting repulsion in misfolded proteins (Amangeldina et al., 2024).

#### 3.3.3 Amino acid usage bias captures both GC content and adaptation to environment

The main driver of amino acid composition is nucleotide bias, followed by temperature (Hickey and Singer, 2004). In our dataset, the amino acid usage bias, measured by the standard deviation of the 20 common amino acid frequencies, is strongly correlated with GC content in GC-rich genomes (GC% *>*= 50%: r = 0.94; GC% *<* 50%: r = -0.49). This trend remains across temperature bins. AAUB correlates moderately with OGT only for AT-rich genomes and in the top 20% of the dataset (r = 0.57) (Supplementary Figure S5).

### 3.4 Phylogenetic signal

Both OGT and the descriptors contain significant evolutionary signal under the Brownian motion model of evolution (Supplementary Table S1). For Archaea, Pagel’s lambda was 0.96 (p *<* 0.001), and for Bacteria 0.92 (p *<* 0.001). Therefore, under a random training-test split, data leakage occurs and an alternative phylogeny-based split should be performed. In reality, the best model is a trade-off between generalization and performance (Cimen et al., 2020). Here, we kept observations of the same genus level together.

### 3.5 Machine learning results

#### 3.5.1 Non-linear models show the most improvement in (hyper)thermophiles

Non-linear models have consistently better cross-validation RMSEs compared to the linear models (RMSE = 6.53°C) overall and specifically in the thermophilic region (Figure 3). The best model based on overall metrics is the Gaussian process model with cross-validation RSME = 5.05 (*±* 0.12) and R^2^ = 0.89 (*±* 0.03), followed by the multilayer perceptron (mlp) model (RMSE = 5.07°C (*±* 0.24) and R^2^ = 0.89 (*±* 0.04)).

**Figure 3:**
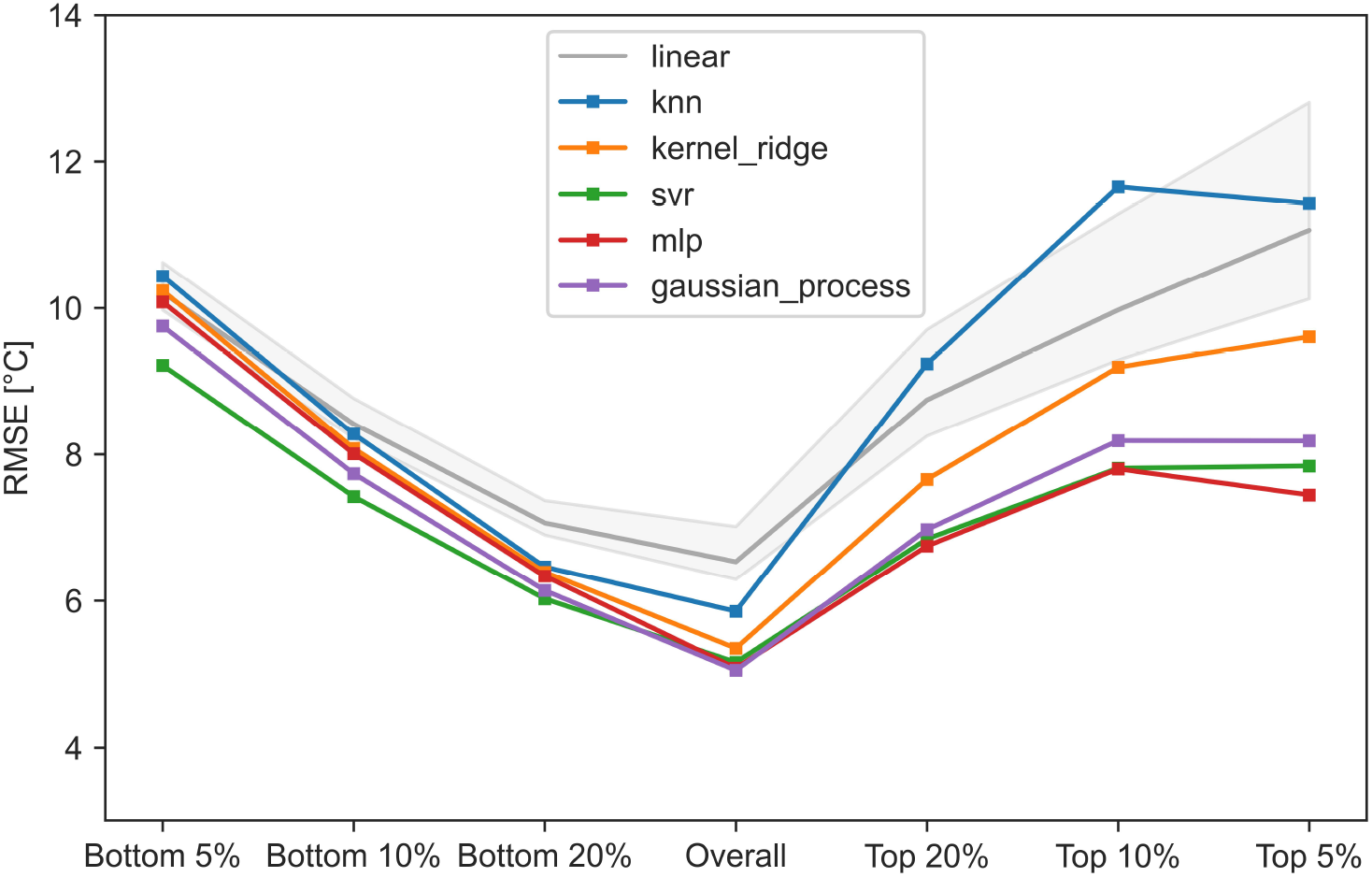
Combined cross-validation RMSE for the individual non-linear models compared to a linear model background (CI = 95%). The percentages indicate the upper and lower 5%, 10%, and 20% quantiles. Improvement over the linear models is stronger at higher temperatures.

#### 3.5.2 Poor fit for psychrophiles

The RMSE of all models increases near the extremes (Figure 3). High R^2^ values are observed in the upper quantiles, indicating that the model performs best at the higher temperature extreme. The reason why all models capture thermophilic trends better than psychrophilic ones is two-fold. First, the dataset consists of only 0.9% (n = 58) psychrophiles compared to 88,9% (n = 5,689) mesophiles and 10.2% (n = 656) (hyper)thermophiles. Given the few psychrophilic observations, the model does not focus enough on accurately predicting these during training. To partially overcome this, we opted for weighted RMSE. Secondly, psychrophiles have weaker signals in their amino acid composition (Yang et al., 2015), making it more challenging for the model to predict them accurately.

### 3.6 Model interpretation

According to the SHAP analysis, the top three models, mlp, svr and gaussian process, share among their most important features the amino acid descriptors B6, MSWHIM3 and F1 (Supplementary Figures S6-S8). Both B6 and F1 have an inverse relation with OGT. The taxonomic domain, included as a dummy variable in the training set, has only a small contribution in comparison.

#### 3.6.1 B6

The amino acid descriptor B6 contains signals of backbone dynamics (Supplementary Table S2), with the lowest value for proline (P) and the highest for glycine (G) (Supplementary Table S3). Low SHAP values for B6 are associated with high OGT, indicating that thermophiles favor more rigid protein structures. As kinetic energy increases with higher temperature, more stability is needed to retain the protein structure which is essential for its function.

#### 3.6.2 MSWHIM3

The amino acid descriptor MSWHIM3 separates the positively charged amino acids lysine (K) and arginine (R) from the other amino acids. Most negative MSWHIM3 values are associated with the negatively charged amino acid aspartate (D) and the polar amino acids serine (S) and threonine (T) (Supplementary Table S3). The relation between charged amino acids and thermal adaptation is complex. Higher fractions of charged residues are typically associated with thermophilic adaptation. The two main reasons are 1) increasing protein thermostability by stabilizing the core and through interaction with the solvent and salt bridges at the protein surface (Hait et al., 2020; Amangeldina et al., 2024) and 2) disfavoring misfolded states through negative design (Amangeldina et al., 2024). In contrast, the fraction of negatively charged, thermolabile aspartate decreases at elevated temperatures (Yang et al., 2015). The MSWHIM3 scores capture this trend, with high values for lysine and arginine and a low value for aspartate (Supplementary Table S3). Moreover, the correlation of the fraction 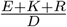 with OGT behaves similarly in GC-rich and GC-poor genomes. This trend is strongest in the top 20% of observations GC% *>*= 50%: r = 0.66; GC% *<* 50%: r = 0.64). Finally, the correlation is more pronounced in Archaea (overall r = 0.77) compared to Bacteria (overall r = 0.41).

#### 3.6.3 F1

The descriptor F1 captures an amino acid’s hydrophobicity and buriability in contrast to the accessible surface area (Supplementary Table S2). Low F1 values exist for all charged amino acids (D, E, K, R) as well as the polar residues asparagine (N) and glutamine (Q) (Supplementary Table S3). Positive F1 values are associated with sulfur-containing amino acids methionine (M) and cysteine (C), and aromatic amino acids tryptophan (W) and phenylalanine (F). In addition, an inconclusive signal is present for bulky, hydrophobic amino acids with low F1 values for valine (V), whereas isoleucine (I) and leucine (L) have positive F1 values. The correlation between F1 and SHAP values is negative, which suggests that thermophiles favor hydrophilic, solvent-accessible amino acids. An increased accessible surface area promotes solvent interaction, which improves thermostability by preventing solvent molecules from entering the protein (Hait et al., 2020). Concerning the sulfur-containing amino acids, no significant correlations were previously found with OGT (Villain et al., 2022; Sauer and Wang, 2019). However, in our dataset, the fraction “CM” is negatively correlated to OGT in the top 20% for AT-rich genomes (r = -0.45), but not in GC-rich genomes (r = -0.09), which may have previously obscured the relation. Finally, aromatic amino acids are typically associated with thermoadaptation as the *π*-*π* interactions contribute to thermostability. However, the overall correlation in our dataset between OGT and the fraction *Y* + *W* + *F* is low (r = 0.15). Furthermore, PCA coloring shows a strong phylogenetic signal (Supplementary Figures S4, S9).

### 3.7 Comparison to existing OGT prediction models

Comparing OGT prediction models directly requires an independent test set, which is challenging due to different training datasets and the need for distantly related species because of the strong phylogenetic signal. Furthermore, comparison of RMSE and R^2^ is unreliable as many models did not include a phylogenetically informed split, resulting in overoptimistic estimates. Still, our mlp model has a test RMSE of 5.49°C and a test R^2^ of 0.836, which is similar to that of the best existing prokaryotic OGT prediction model (RMSE = 5.17, R^2^ = 0.835; Sauer and Wang 2019), while including a phylogenetic split on the genus level.

## 4 Conclusion

In this work we constructed a new (optimal) growth dataset containing OGT values for 10,582 unique TaxIds from all three domains of life, although prokary-otes are heavily overrepresented. For the 6,401 TaxIds with available genomes we built OGT prediction models. The best model was the multilayer perceptron with a cross-validation RMSE of 5.07°C (±0.24) and an R^2^ equal to 0.89 (±0.04). This performance is similar to the state-of-the-art general prokaryotic OGT prediction model of Sauer and Wang (2019), even though we included a phylogenetic split on the genus level to reduce data leakage between training and test set. The most important predictors were related to backbone flexibility (B6), charged residues (MSWHIM3), and surface accessibility (F1). Moreover, fractions of the sulfur-containing amino acids cysteine and methionine were significantly decreased in (hyper)thermophiles compared to mesophiles, but only significantly in AT-rich genomes.

## 5 Data availability

The growth temperature dataset is available in supplementary data and hosted on Zenodo (doi: 10.5281/zenodo.15282730). OGTFinder is available on GitHub and archived on Zenodo (doi: 10.5281/zenodo.15282984).

## 7 Acknowledgments

This work was supported by KU Leuven project C1 ‘ACES’ [C16/20/001]; and VLAIO project ‘ENZYMARES’ [HBC.2021.0076]. J.F holds a PhD fellowship strategic basic research (grant number: 1SE4625N) of the Research Foundation - Flanders (FWO, Belgium). The resources and services used in this work were provided by the VSC (Flemish Supercomputer Center), funded by the Research Foundation - Flanders (FWO) and the Flemish Government.

## Footnotes

1 https://www.ccug.se/

3 https://www.uv.es/uvweb/spanish-type-culture-collection/en/spanish-type-culture-collection-1285872233521.html

3 https://mcc.nies.go.jp/

