## Supplementary for "Machine learning for optimal growth temperature prediction of prokaryotes using amino acid descriptors"

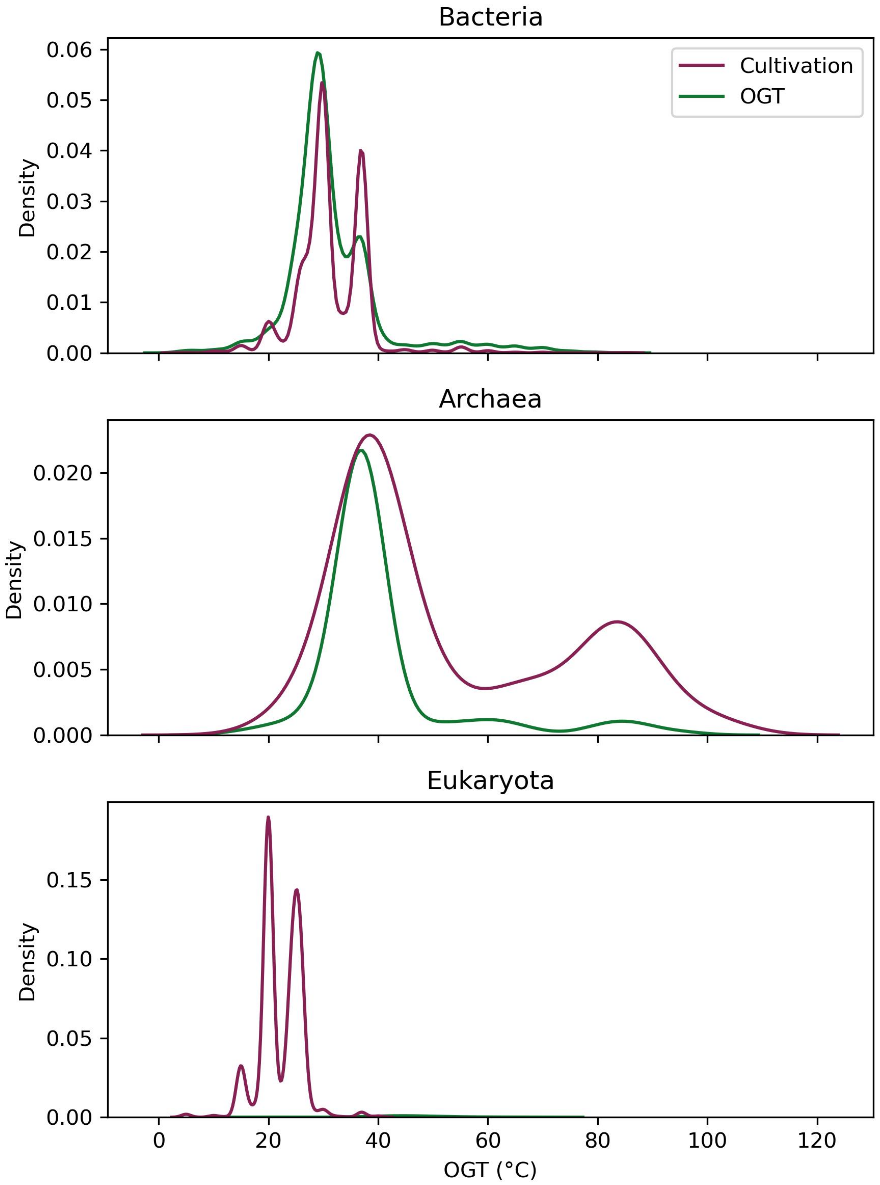


Figure S1: Kernel density estimations on cultivation and optimal growth temperatures for the three domains of life.

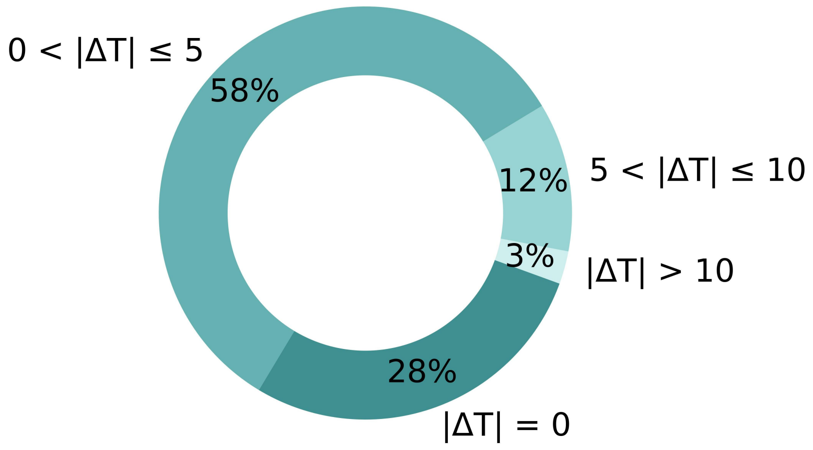


Figure S2: Distribution of the absolute deviations between median cultivation temperature and median OGT for 2,598 species.

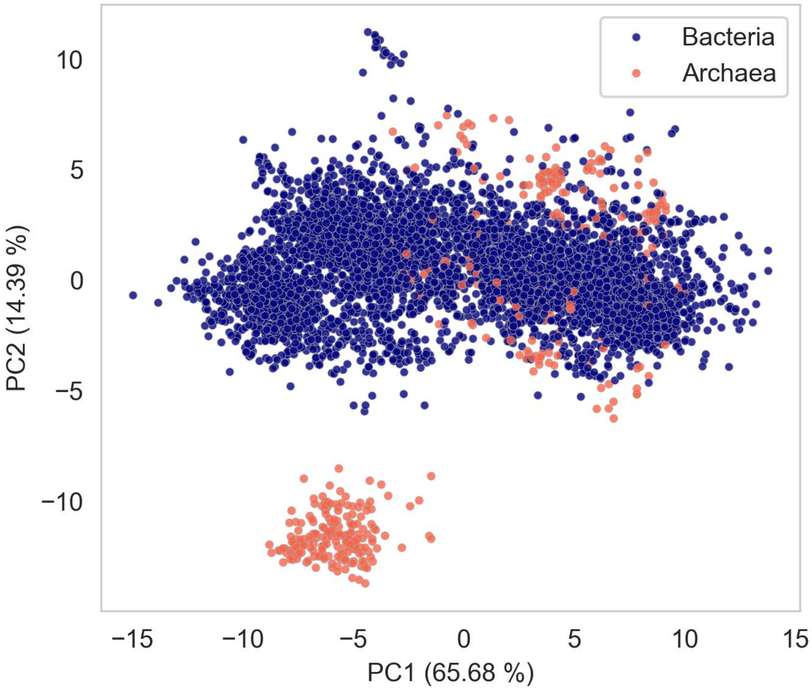


Figure S3: First two principal components (PCs) for the 6,401 taxids in the OGT dataset, colored by the taxonomic domain.

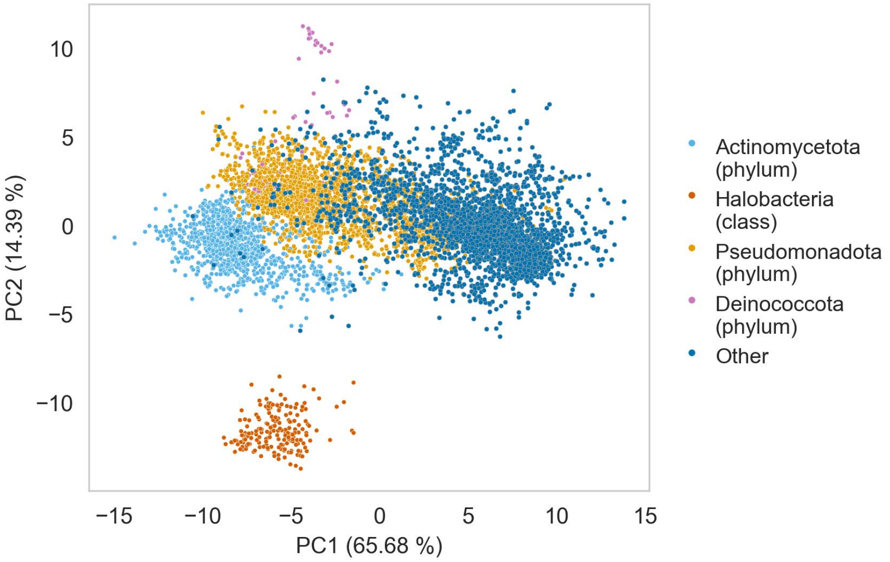


Figure S4: Some (partially) isolated taxa in the plot of the first two principal components (PCs) for the 6,401 taxids in the OGT dataset.

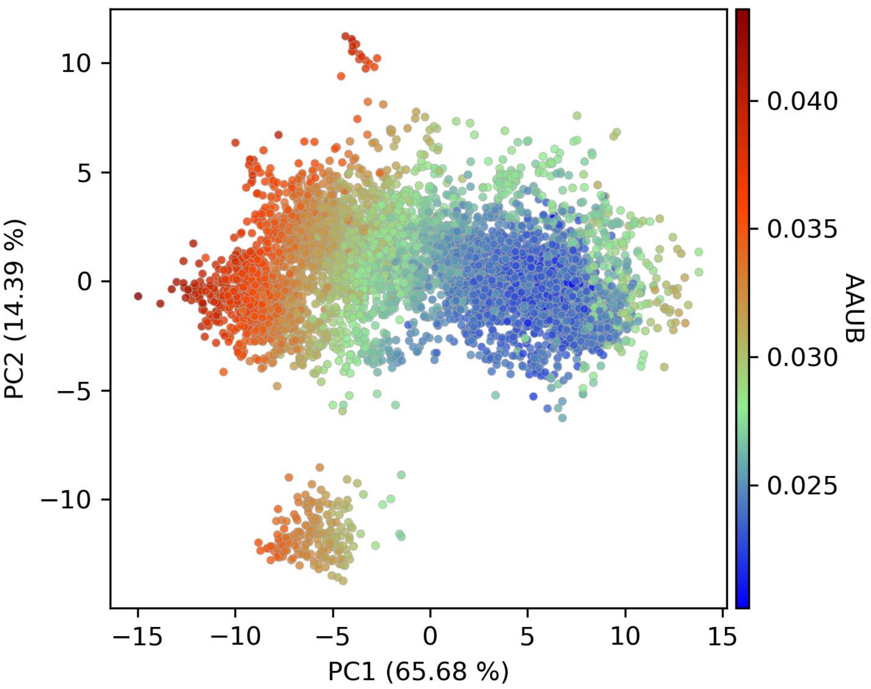


Figure S5: First two principal components (PCs) for the 6,401 taxids in the OGT dataset, colored by the sample standard deviation over the 20 common amino acid frequencies (= amino acid usage bias, AAUB). AAUB captures both GC-content and environmental factors.

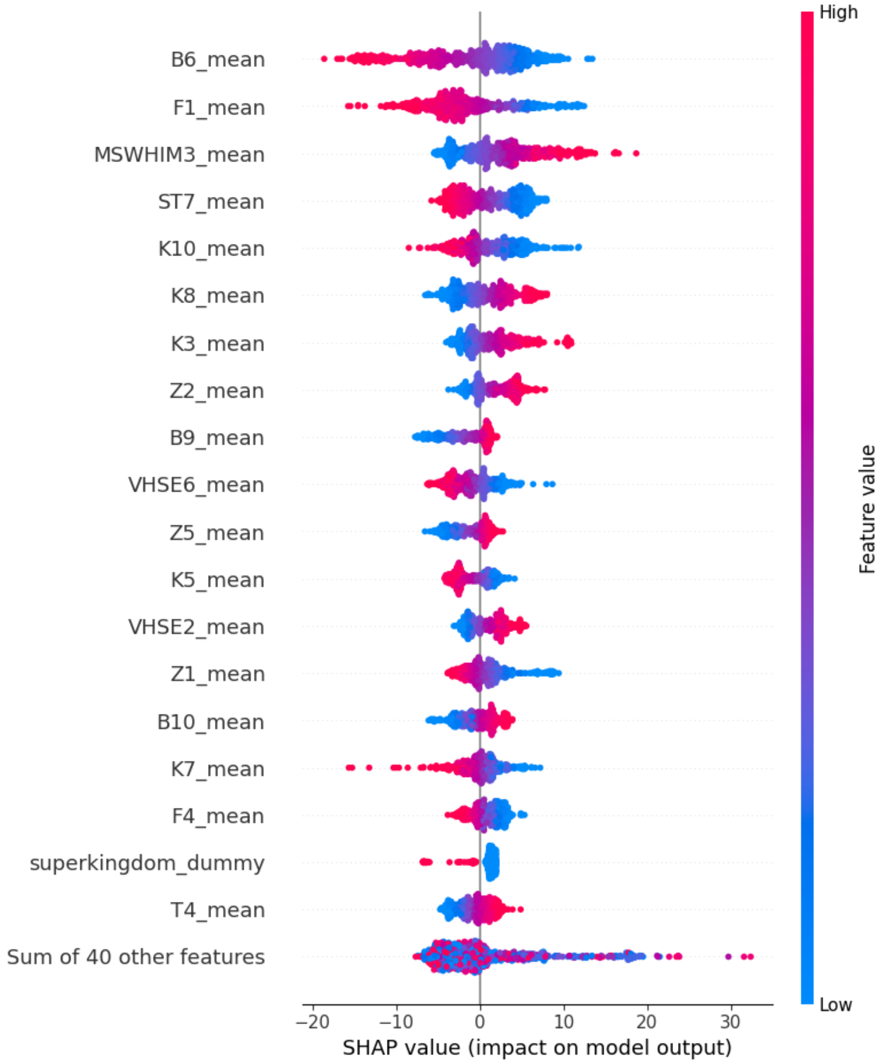


Figure S6: Beeswarm summary plot of the top 20 features from SHAP analysis on the multi-layer perceptron regression model with test observations.

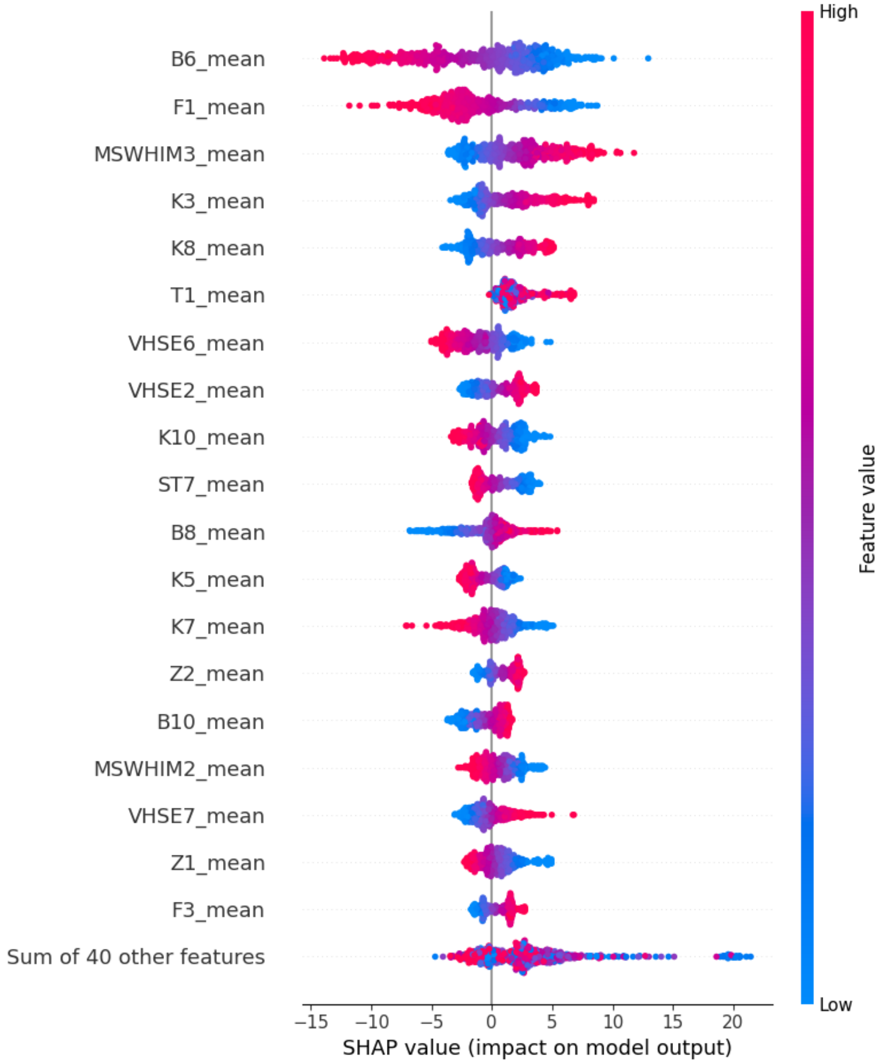


Figure S7: Beeswarm summary plot of the top 20 features from the SHAP analysis on support vector regression model with test observations.

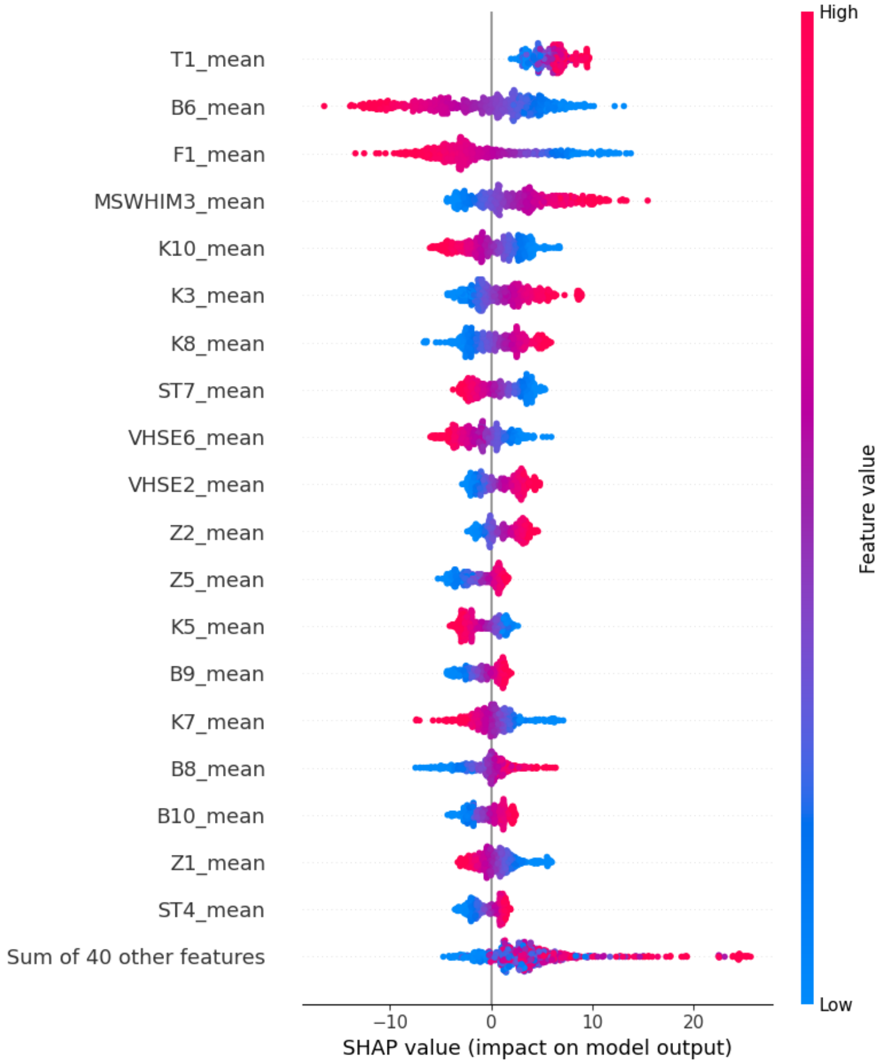


Figure S8: Beeswarm summmary plot of the top 20 features from the SHAP analysis on gaussian process regression model with test observations.

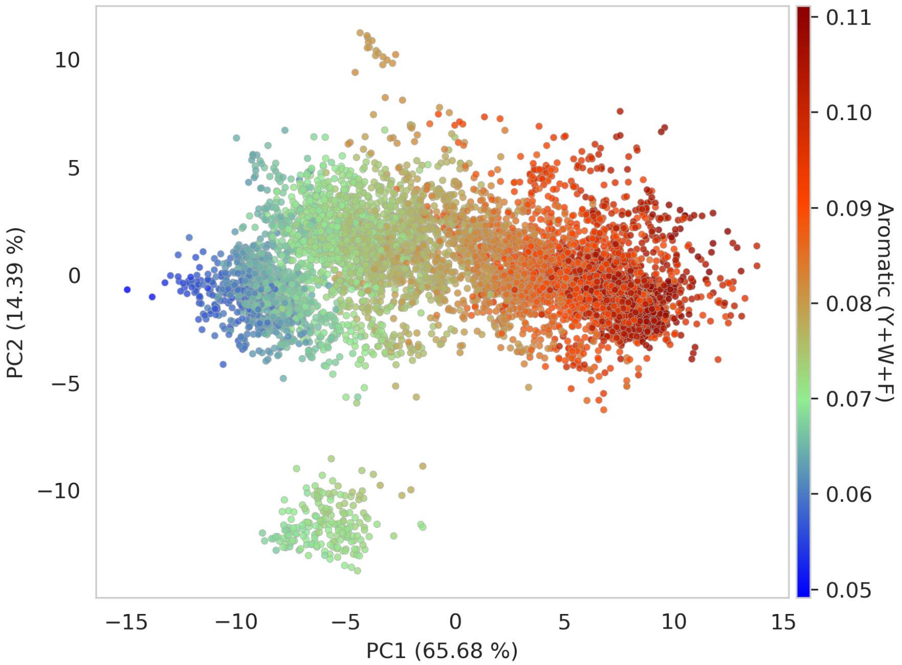


Figure S9: First two principal components (PCs) for the 6,401 taxids in the OGT dataset, colored by aromatic amino acid fraction (Y+W+F). The aromatic fraction roughly follows AT-content/phylogeny.

### Supplementary Tables

Table S1: Pagel's $\lambda$ values for predictors (averaged amino acid descriptors) and response (median OGT) in Bacteria and Archaea, calculated using a time-scaled tree from TimeTree, with zero-length branches removed. All values are significant (Benjamini-Hochberg corrected p-value $< 0.001$).

| Predictor | Pagel's $\lambda$ | |
|---|---|---|
| | Bacteria (n = 3,829) | Archaea (n = 298) |
| OGT (median) | 0.92 | 0.96 |
| K1 | 0.96 | 0.99 |
| K2 | 0.98 | 0.98 |
| K3 | 0.98 | 0.99 |
| K4 | 0.98 | 1.00 |
| K5 | 0.93 | 0.98 |
| K6 | 0.97 | 0.99 |
| K7 | 0.98 | 0.99 |
| K8 | 0.92 | 0.99 |
| K9 | 0.97 | 0.99 |
| K10 | 0.98 | 1.00 |
| T1 | 0.97 | 0.98 |
| T2 | 0.98 | 0.99 |
| T3 | 0.95 | 0.98 |
| T4 | 0.97 | 0.99 |
| T5 | 0.94 | 0.98 |
| ST1 | 0.99 | 0.99 |
| ST2 | 0.95 | 0.97 |
| ST3 | 0.98 | 1.00 |
| ST4 | 0.94 | 0.99 |
| ST5 | 0.91 | 0.99 |
| ST6 | 0.98 | 0.99 |
| ST7 | 0.96 | 0.99 |
| ST8 | 0.96 | 0.97 |
| B1 | 0.98 | 0.99 |
| B2 | 0.95 | 0.99 |
| B3 | 0.96 | 1.00 |
| B4 | 0.97 | 0.99 |
| B5 | 0.98 | 1.00 |

| Predictor | Pagel's $\lambda$ | |
| --- | --- | --- |
| | Bacteria (n = 3,829) | Archaea (n = 298) |
| B6 | 0.98 | 0.99 |
| B7 | 0.97 | 0.97 |
| B8 | 0.95 | 0.98 |
| B9 | 0.97 | 0.99 |
| B10 | 0.98 | 0.98 |
| Z1 | 0.95 | 0.96 |
| Z2 | 0.98 | 0.99 |
| Z3 | 0.98 | 0.99 |
| Z4 | 0.97 | 0.99 |
| Z5 | 0.98 | 0.99 |
| MSWHIM1 | 0.98 | 1.00 |
| MSWHIM2 | 0.97 | 0.99 |
| MSWHIM3 | 0.98 | 0.98 |
| VHSE1 | 0.98 | 0.98 |
| VHSE2 | 0.97 | 1.00 |
| VHSE3 | 0.96 | 0.96 |
| VHSE4 | 0.98 | 0.99 |
| VHSE5 | 0.95 | 0.98 |
| VHSE6 | 0.98 | 0.99 |
| VHSE7 | 0.98 | 0.99 |
| VHSE8 | 0.96 | 0.98 |
| F1 | 0.96 | 0.99 |
| F2 | 0.96 | 0.98 |
| F3 | 0.94 | 0.99 |
| F4 | 0.97 | 0.99 |
| F5 | 0.96 | 0.99 |
| F6 | 0.98 | 0.98 |
| PP1 | 0.97 | 0.99 |
| PP2 | 0.96 | 0.99 |
| PP3 | 0.97 | 0.99 |

Table S2: Top Pearson correlations of AA indices with key descriptors. Categories were derived from AAontology.

| | Top hits | | |
|---|---|---|---|
| **Descriptor** | **Index** | **r** | **Category** |
| B6 | ROBB760107 | -0.76 | Conformation, Coil (C-term), Coil (C-terminal), "Information measure for extended without H-bond" |
| | FINA910102 | -0.69 | Conformation, Linker (>14 AA), $\alpha$-helix (C-term, out), "Helix initiation parameter at position i,i+1,i+2" |
| | ONEK900101 | 0.65 | Others, Unclassified (Others), $\Delta$G values in peptides, "Delta G values for the peptides extrapolated to 0 M urea" |
| | BUNA790101 | 0.71 | Structure-Activity, Backbone-dynamics (-NH), $\alpha$-NH chemical shifts (backbone-dynamics), "alpha-NH chemical shifts" |
| MSWHIM3 | QIAN880125 | -0.67 | Conformation, $\beta$-sheet (C-term), $\beta$-sheet (C-terminal), "Weights for beta-sheet at the window position of 5" |
| | QIAN880126 | -0.64 | Conformation, $\beta$-sheet (C-term), $\beta$-sheet (C-terminal), "Weights for beta-sheet at the window position of 6" |
| | FINA770101 | 0.61 | Structure-Activity, Stability (helix-coil), Stability (helix-coil), "Helix-coil equilibrium constant" |
| | RICJ880112 | 0.66 | Conformation, $\alpha$-helix, $\alpha$-helix,"Relative preference value at C3" |
| F1 | FAUJ830101 | 0.85 | Polarity, Hydrophobicity, Hydrophobicity, "Hydrophobic parameter $\pi$" |
| | MEIH800103 | 0.84 | Polarity, Amphiphilicity ($\alpha$-helix), Side chain angle (phi), "Average side chain orientation angle" |
| | BIOV880101 | 0.83 | ASA/Volume, Buried, Buriability, "Information value for accessibility; average fraction 35%" |
| | GRAR740102 | -0.84 | Polarity, Hydrophilicity, Polarity (hydrophilicity), "Polarity" |

Table S3: Descriptor values per amino acid for the key descriptors MSWHIM3, F1 and B6.

| Amino acid | Abbreviation | MS-WHIM3 | F1 | B6 |
|---|---|---|---|---|
| Leu | L | -0.16 | 1.200 | 0.34 |
| Ile | I | -0.25 | 1.524 | 0.28 |
| Met | M | -0.32 | 0.886 | 0.37 |
| Pro | P | -0.60 | -0.407 | -2.02 |
| Asn | N | -0.66 | -1.009 | 0.83 |
| Gln | Q | -0.30 | -0.880 | -0.08 |
| Lys | K | 0.60 | -1.387 | 0.10 |
| Arg | R | 1.00 | -1.229 | 0.20 |
| Gly | G | -0.75 | -0.205 | 1.19 |
| Ala | A | -0.62 | 0.207 | 0.19 |
| Cys | C | -0.27 | 0.997 | -1.05 |
| Ser | S | -1.00 | -0.495 | 0.54 |
| Val | V | -0.58 | -1.332 | 0.16 |
| Thr | T | -0.89 | -0.032 | 0.38 |
| Asp | D | -0.96 | -1.298 | 0.01 |
| Glu | E | -0.04 | -1.349 | -0.08 |
| His | H | -0.78 | -0.270 | -0.79 |
| Phe | F | -0.34 | 1.247 | 0.29 |
| Trp | W | -0.47 | 0.844 | 0.24 |
| Tyr | Y | -0.16 | 0.329 | -0.48 |
